## Supplemental Material for "The contribution of abortive infection to preventing populations of *Lactococcus lactis* from succumbing to infections with bacteriophage"

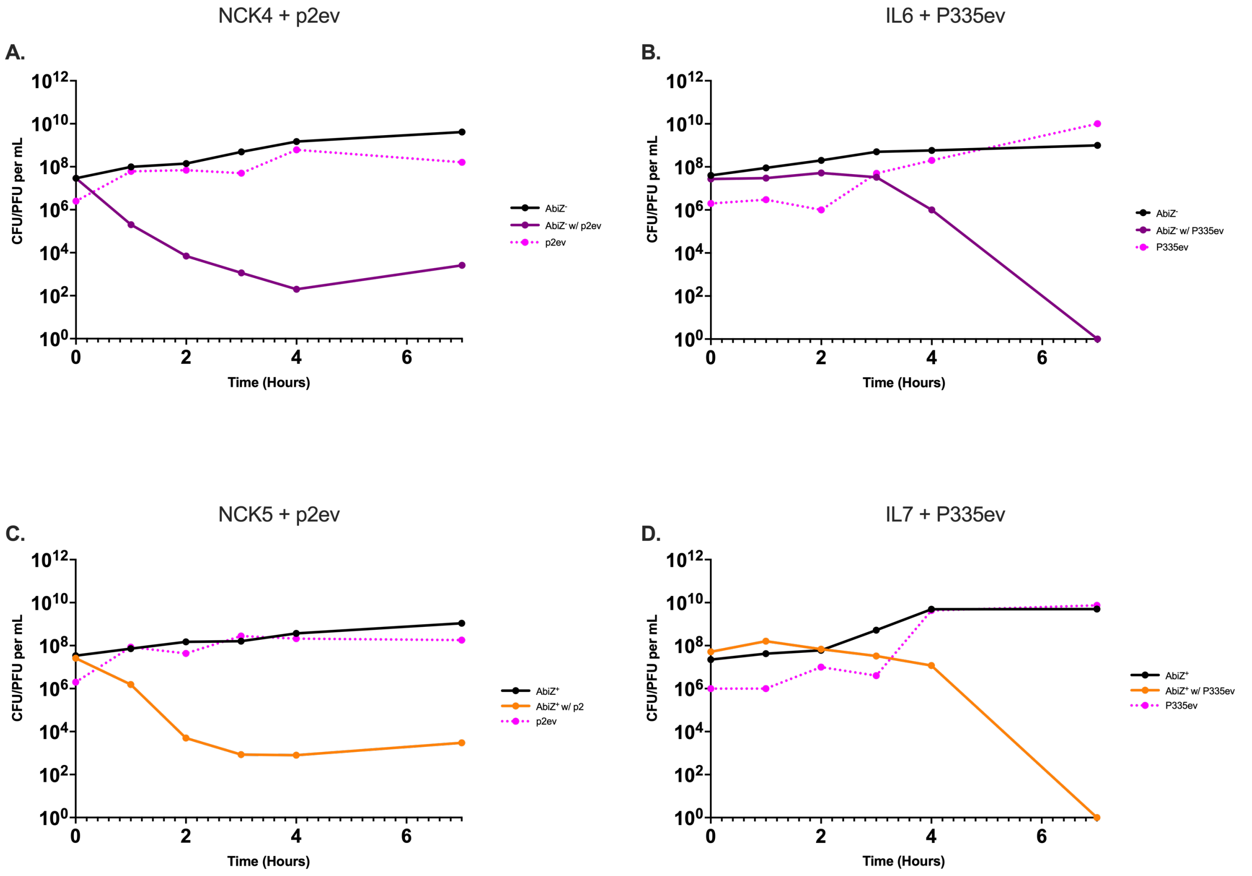


Figure S1. **Short-term population dynamics of L. lactis with evolved phage.** Densities of bacteria and respective ancestral and evolved phage over the course of 7 hours post infection. Solid black line represents abiZ- or abiZ+ controls grown without phage present. A- AbiZ- cells (purple) with phage P2ev (dashed pink). B- AbiZ- cells (purple) with phage P335ev (dashed pink). C- AbiZ+ cells (orange) with phage P2ev D- AbiZ+ cells (orange) with P335ev.

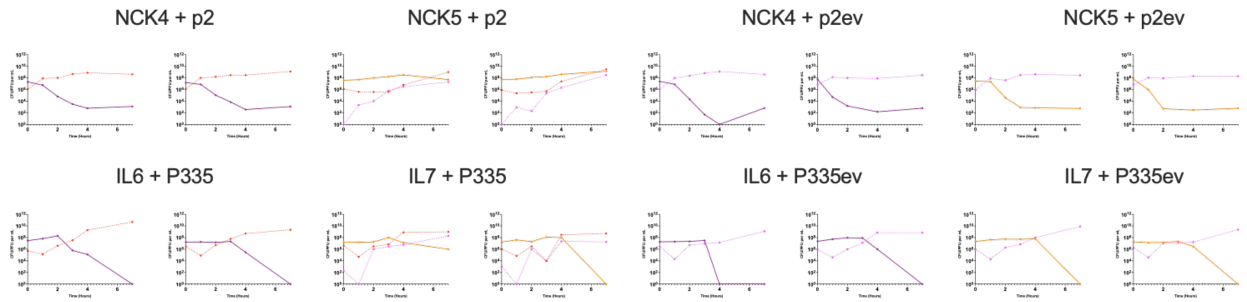


**Figure S2. Short term (7 hour) experiment biological replicas.** $AbiZ^-$ (NCK4/IL6) bacteria (purple) total phage p2/P335 (dashed red), $AbiZ^+$ (NCK5/IL7) cells (orange), phage p2ev/P335ev (dashed pink)

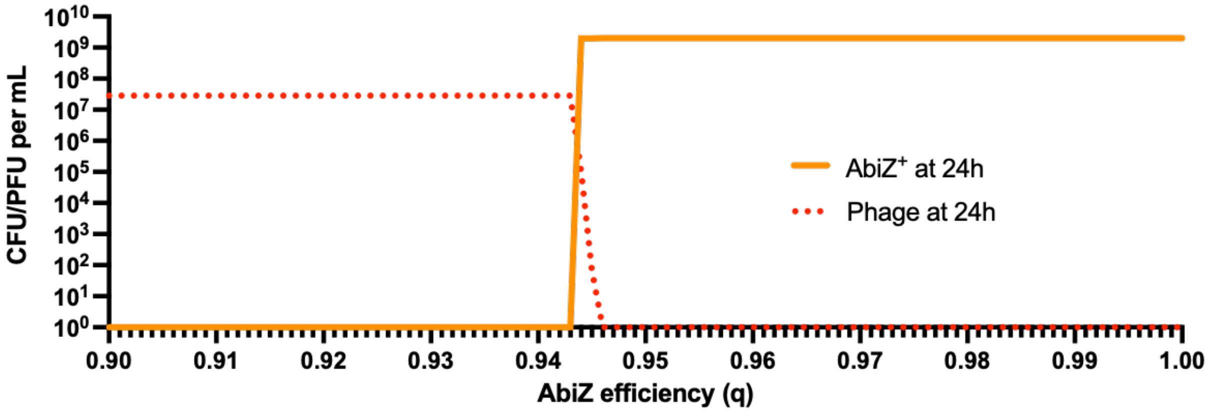


**Figure S3. Computer simulation results for the effect of Abi efficiency as values of q change**.

Plotted are the 24-hour densities of phage and $AbiZ^+$ bacteria for varying values of q. The parameters used are the same as those in Table 1 without the transition to and from resistant.

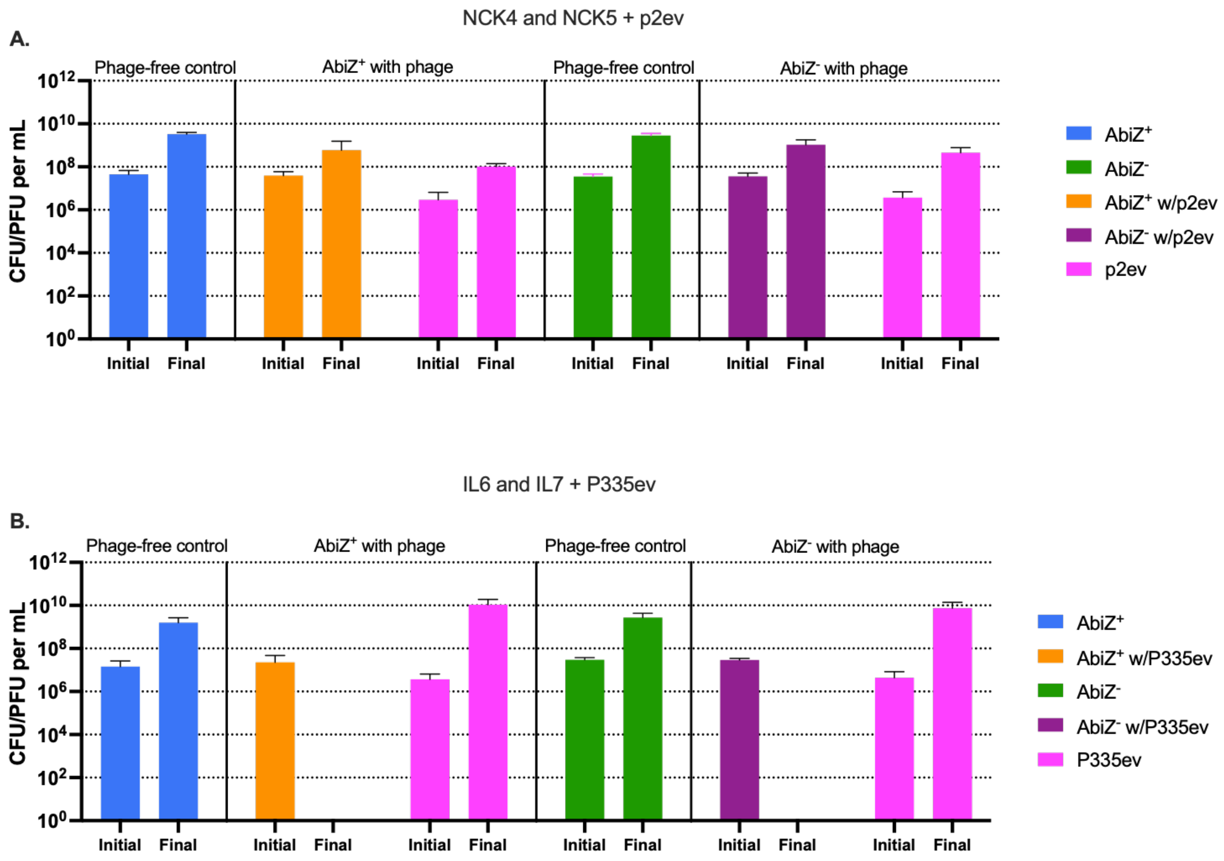


**Figure S4. Conditions for abortive infection protection against p2ev or P335ev following 24 hours in liquid culture**. Bars represent mean initial (Time=0hr) and final (Time=24hr) colony or plaque forming units per mL of 3 biological-replicas and error bars represent ± SD. A- P2ev. B- P335ev. Blue bars represent cells lacking Abi not confronted by phage. Green bars represent cells lacking Abi which are not confronted by phage. Orange and purple bars represent cells with or without Abi respectively cocultured with phage. p2ev/P335ev is represented by pink bars.

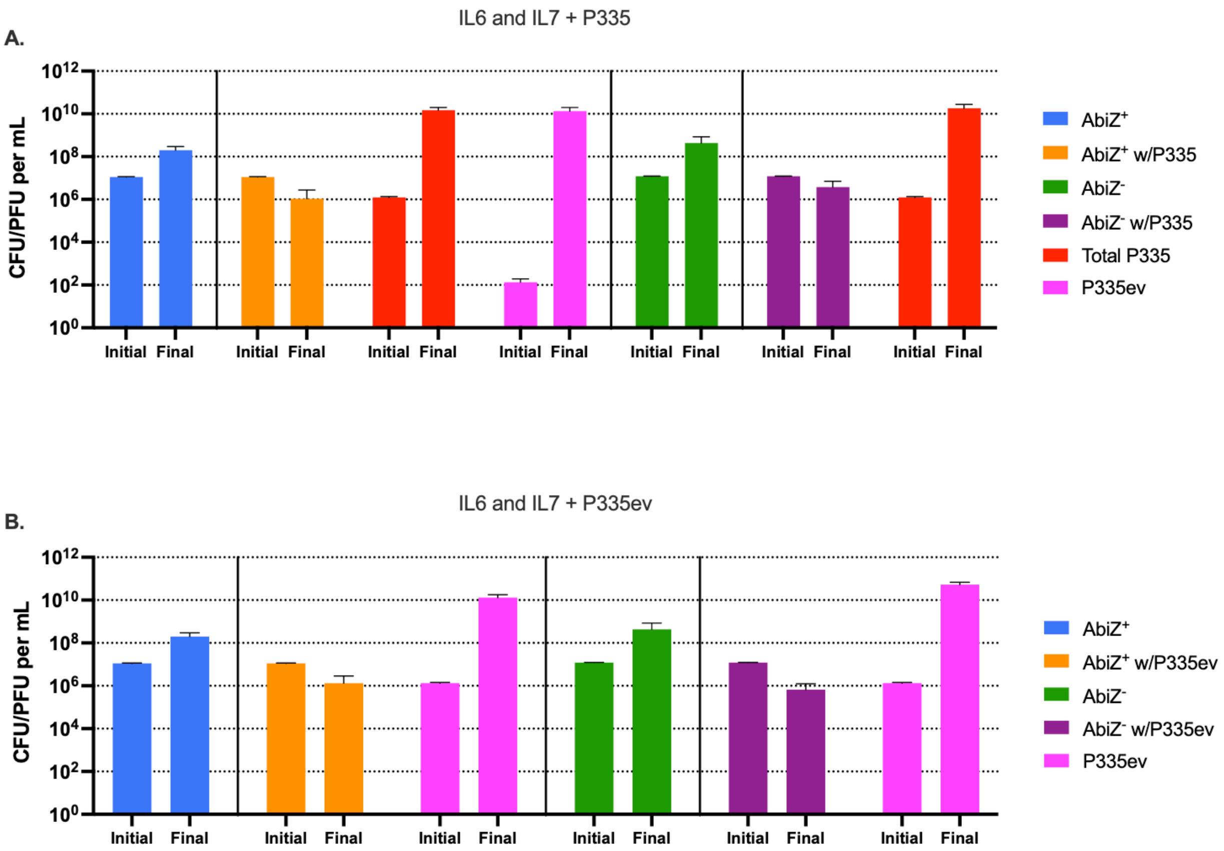


**Figure S5. Conditions for abortive infection protection against P335 or P335ev following 96 hours in liquid culture**. Bars represent mean initial (Time=0hr) and final (Time=24hr) colony or plaque forming units per mL of 3 biological-replicas and error bars represent ± SD. A- P335. B- P335ev. Blue bars represent cells lacking Abi not confronted by phage. Green bars represent cells lacking abi which are not confronted by phage. Orange and purple bars represent cells with or without Abi respectively cocultured with phage. P335 is represented by red bars and P335ev is represented by pink bars.

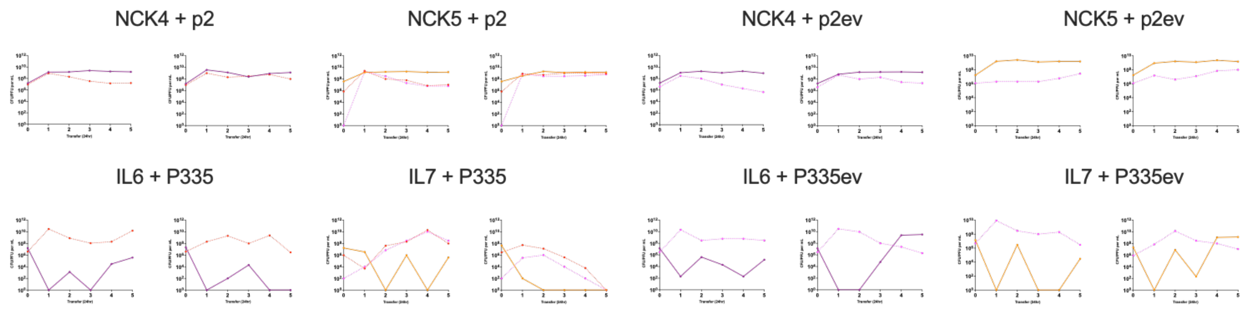


**Figure S6. Biological replicas of serial transfer experiments**. AbiZ- (NCK4/IL6) bacteria (purple) total phage p2/P335 (dashed red), AbiZ+ (NCK5/IL7) cells (orange), phage p2ev/P335ev (dashed pink).

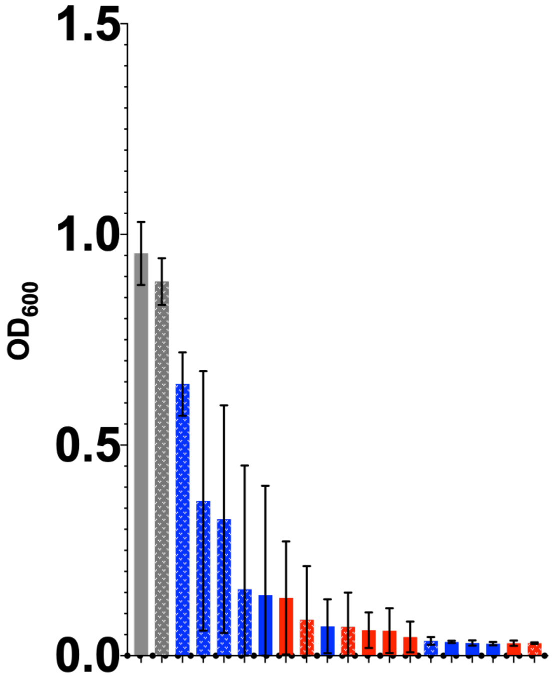


**Figure S7. MaxOD after 24 hours of isolates from IL6/IL7 serial transfer experiments days 2-5 grown with 1e7 P335ev.** Bars show mean of 5 technical replicas with error bars ± SEM. Solid bars represent $AbiZ^-$ IL6 and dotted bars represent $AbiZ^+$ IL7. Blue bars are isolates which appeared resistant by spot testing, red bars are isolates which appeared sensitive by spot testing, grey bars are IL6 and IL7 grown without phage.

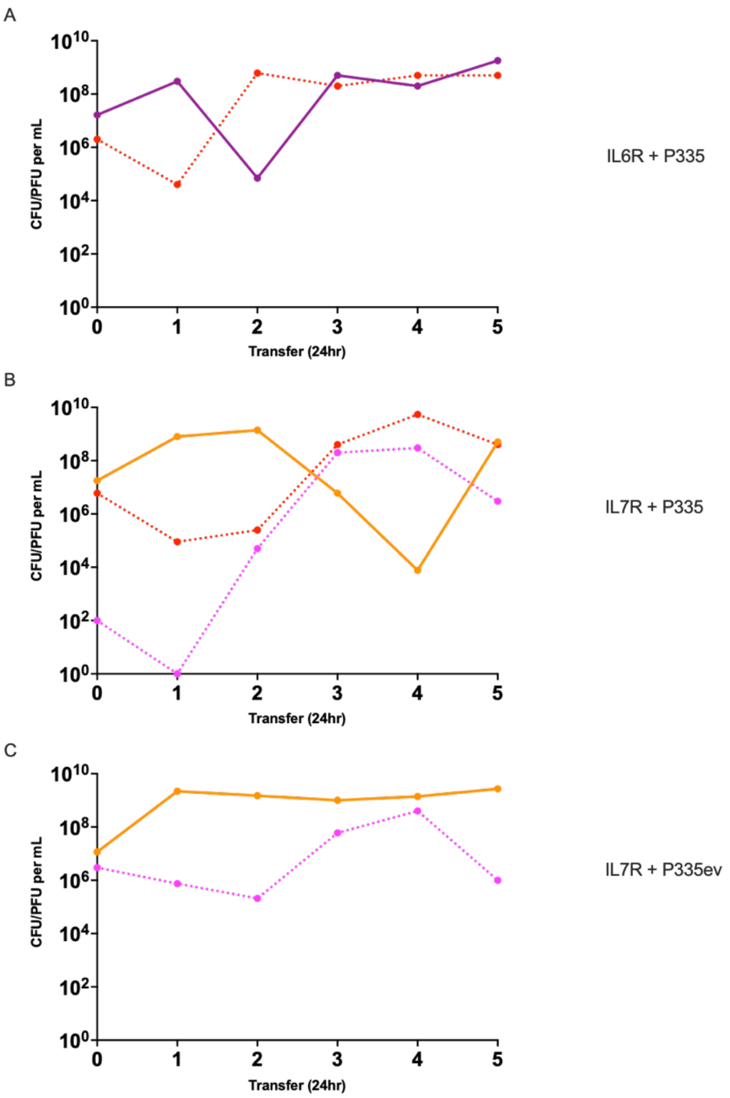


**Figure S8. Serial transfers of 96-hour IL6 and IL7 resistant mutants with P335 and P335ev.** Purple line represents AbiZ$^{-}$ IL6 bacteria, orange line represents AbiZ$^{+}$ IL7 bacteria, red dotted line represents Total P335 phage, and pink dotted line represents P335ev.

**Table S1. Point mutations in phages p2 and P335ev.** p2ev' is phage isolated after 7 hours in liquid with AbiZ+, p2ev is after 24 hours in the same experimental conditions.

| Region | Phage | | |
|---|---|---|---|
| | p2ev' | p2ev | P335ev |
| Major Capsid Protein | Gln178Lys | Gln178Lys | Thr268Lys |
| Receptor Binding Protein | | Met165Thr | |

**Table S2. Proportion of resistant cells recovered following 24 hours liquid experiments by spot testing.**

| *L. lactis* + Phage | Figure | Proportion of Resistant Bacteria |
|---|---|---|
| NCK5 + p2 | 4A | 0.2 |
| NCK4 + p2 | 4A | 0.9 |
| NCK5 + p2ev | S4A | 1 |
| NCK4 + p2ev | S4A | 0.9 |
| IL7 + P335 | 4B | 0 |
| IL6 + P335 | 4B | No recovery |
| IL7 + P335ev | S4B | No recovery |
| IL6 + P335ev | S4B | No recovery |

**Table S3. Proportion of resistant cells recovered following 96 hours liquid experiments by spot testing.**

| *L. lactis* + Phage | Figure | Proportion of Resistant Bacteria |
|---|---|---|
| IL7 + P335 | S5A | 1 |
| IL6 + P335 | S5A | 1 |
| [illegible]L7 + P335ev | S5B | 1 |
| IL6 + P335ev | S5B | 1 |

**Table S4. Proportion of resistant cells recovered during serial transfers by spot testing for NCK4/NCK5 and p2.**

| ***L. lactis*** **+ Phage** | **Figure** | **Proportion of Resistant Bacteria (Tfr. 1, 2, 3, 4, 5)** |
|---|---|---|
| NCK4 + p2 | 5A | 1, 1, 1, 1, 1 |
| NCK5 + p2 | 5C | 0 , 0.33, 1, 1, 1 |
| NCK4 + p2ev | 5B | 1,1,1,1,1 |
| NCK5 + p2ev | 5D | 1,1,1,1,1 |
